# Subicular Plateaus Signal Reward Locations during Goal-Directed Behavior

**DOI:** 10.64898/2026.02.16.706209

**Authors:** Aanchal Bhatia, Mohamed Adel, Christine Grienberger

## Abstract

The hippocampus is essential for spatial learning, yet how its principal output structure, the subiculum, encodes behaviorally relevant information during navigation remains poorly understood. Because subicular neurons exhibit prominent burst firing, elucidating how this bursting is generated during behavior is critical. Using in vivo whole-cell recordings from dorsal subiculum neurons in head-fixed mice performing a goal-directed navigation task, we show that many bursts correspond to plateau events consisting of a series of action potentials riding on a large (∼20-30 mV) sustained subthreshold depolarization. Subicular plateaus were synaptic in origin and required NMDA receptor activation. Organizing plateaus by the membrane potential preceding their onset revealed distinct behavioral associations. Plateaus initiated from relatively hyperpolarized membrane potentials preferentially clustered near reward locations, whereas plateaus initiated from more depolarized membrane potentials were more spatially distributed and showed weaker reward specificity. Reward-related clustering required a fixed, learned reward location and dynamically rearranged during learning of a new reward location. Together, our findings identify subicular plateaus as a prominent cellular substrate of hippocampal bursting output and demonstrate that intracellular membrane potential state organizes the behavioral associations of subicular plateaus during goal-directed learning.

## Introduction

The subiculum, a principal output region of the hippocampal formation^1^, is essential for goal-directed navigation and spatial learning^2–4^ and projects to various cortical and subcortical targets^5–7^. Subicular pyramidal neurons receive distinct types of synaptic input. Dense projections from hippocampal area CA1 convey spatial signals, while input from the entorhinal cortex has been linked to contextual and motivational variables^8^. In addition, subicular neurons form dense recurrent connections^9^. Together, this combination of inputs and local connectivity may enable subicular neurons to conjunctively encode multiple behavioral and environmental variables, including spatial location and movement-related variables^6,10–13^.

A prominent physiological feature of subicular pyramidal neurons is their propensity to fire bursts of action potentials (APs)^14,15^. Bursts are known to contribute significantly to information processing in central neurons^16^. While single APs may fail to elicit postsynaptic responses or evoke only weak downstream activation, bursts provide a more reliable means of signal transmission^17,18^ and have been proposed to carry distinct instructive or modulatory signals within hierarchical circuits^19^. Studies have long emphasized bursting as a defining feature of subicular neurons, leading to their classification into regular-spiking and intrinsically bursting cell types based on responses to depolarizing current injection^9,14,15,20–24^. These two neuron types differ in morphology, long-range and local connectivity, and their functional properties^9,25,26^. For instance, the propensity for intrinsic bursting has been shown in acute slice experiments to depend on an afterdepolarization mediated by voltage-gated Ca^2+^ conductances^27^ and to vary across the subiculum, with intrinsically bursting neurons being more prevalent in the distal than in the proximal region^28^.

One candidate mechanism linking burst firing to neuronal information processing in intact circuits is the occurrence of so-called complex spike bursts, which consist of a burst of high-frequency APs riding on a large, plateau-like subthreshold membrane potential (V_m_) depolarization^29–31^ corresponding to a dendritic plateau potential. In cortical and other hippocampal regions, plateau potentials are large (∼20-30 mV) dendritically initiated, non-linear voltage events that depend on synaptic and voltage-gated conductances^32–34^. These events have previously been proposed to signal behaviorally relevant conditions by transiently elevating neuronal output, for example, within the place field of a hippocampal place cell^30,35,36^, and to induce synaptic plasticity^37–39^. Whether similar events occur in the subiculum during active behavior and, if so, whether their expression is structured by behavioral variables remains unknown.

To address this question, we used *in vivo* whole-cell recordings from dorsal subiculum neurons in head-fixed mice performing a goal-directed navigation task. This approach enabled unambiguous identification of plateau-associated complex spike bursts (hereafter referred to as “plateaus”) based on their underlying subthreshold depolarization, a signature inaccessible to extracellular recordings. We find that plateaus are a common feature of subicular pyramidal neuron activity during navigation and depend on NMDA receptor-mediated synaptic transmission. Analysis of plateaus in relation to the pre-plateau V_m_ revealed systematic differences in their behavioral associations: plateaus occurring at relatively hyperpolarized V_m_ values preferentially clustered near reward locations, whereas those occurring at more depolarized V_m_ were more broadly distributed. Together, these results suggest that a neuron’s subthreshold V_m_, driven, among other factors, by synaptic activity, shapes how behaviorally relevant signals are transmitted from the hippocampus to its downstream targets.

## Results

### Whole-cell recordings of dorsal subiculum neurons during goal-directed behavior

To link subthreshold V_m_ dynamics to AP output in a major hippocampal output region during goal-directed behavior, we performed *in vivo* whole-cell recordings from dorsal subiculum neurons in head-fixed mice running on a linear track treadmill for reward at a fixed location (Fig. 1A). Using the blind patch-clamp technique, we recorded from both pyramidal neurons and interneurons; however, interneurons were manually identified based on spike waveform properties and subthreshold V_m_ dynamics and excluded from all further analyses. Accordingly, all results presented in this study are derived from subiculum pyramidal neurons, as confirmed by post hoc biocytin labeling (Fig. 1B), and recovered neurons spanned most of the proximal-distal axis (Fig. 1C). Mean running AP firing rates of subiculum neurons increased from proximal to distal (Fig. 1C), consistent with prior extracellular recordings^6,40^ and were substantially higher than those reported for CA3 or CA1 neurons^41–43^. We further observed that subiculum neurons fired single APs, regular bursts (two or more APs with an inter-spike interval <10 ms and without any large underlying depolarization), and complex spike bursts or “plateaus” (Fig. 1D–F), which were similar to events previously described in hippocampal^30,31,34,36,44^ and cortical^39,45,46^ regions (Fig. 1D-K). Next, we examined baseline V_m_ (zero current through the patch pipette) and firing properties (Fig. 1G–J, n=54 neurons). Mean baseline V_m_ (mean±SEM, running: −54±0.4 mV; standing: −55±0.3 mV) and firing rate (running: 12.0±1.4 Hz; standing: 10.8±1.1 Hz) were similar between running and standing periods, whereas burst rate was modestly higher during running (Fig. 1G–I; mean±SEM, running: 1.7± 0.3 Hz; standing: 1.2±0.2 Hz). The relationship between V_m_ and firing rate, however, was comparable during running and standing (Fig. 1J), implying that increased burst occurrence during running could not be attributed solely to global depolarization or changes in AP gain.

**Figure 1:**
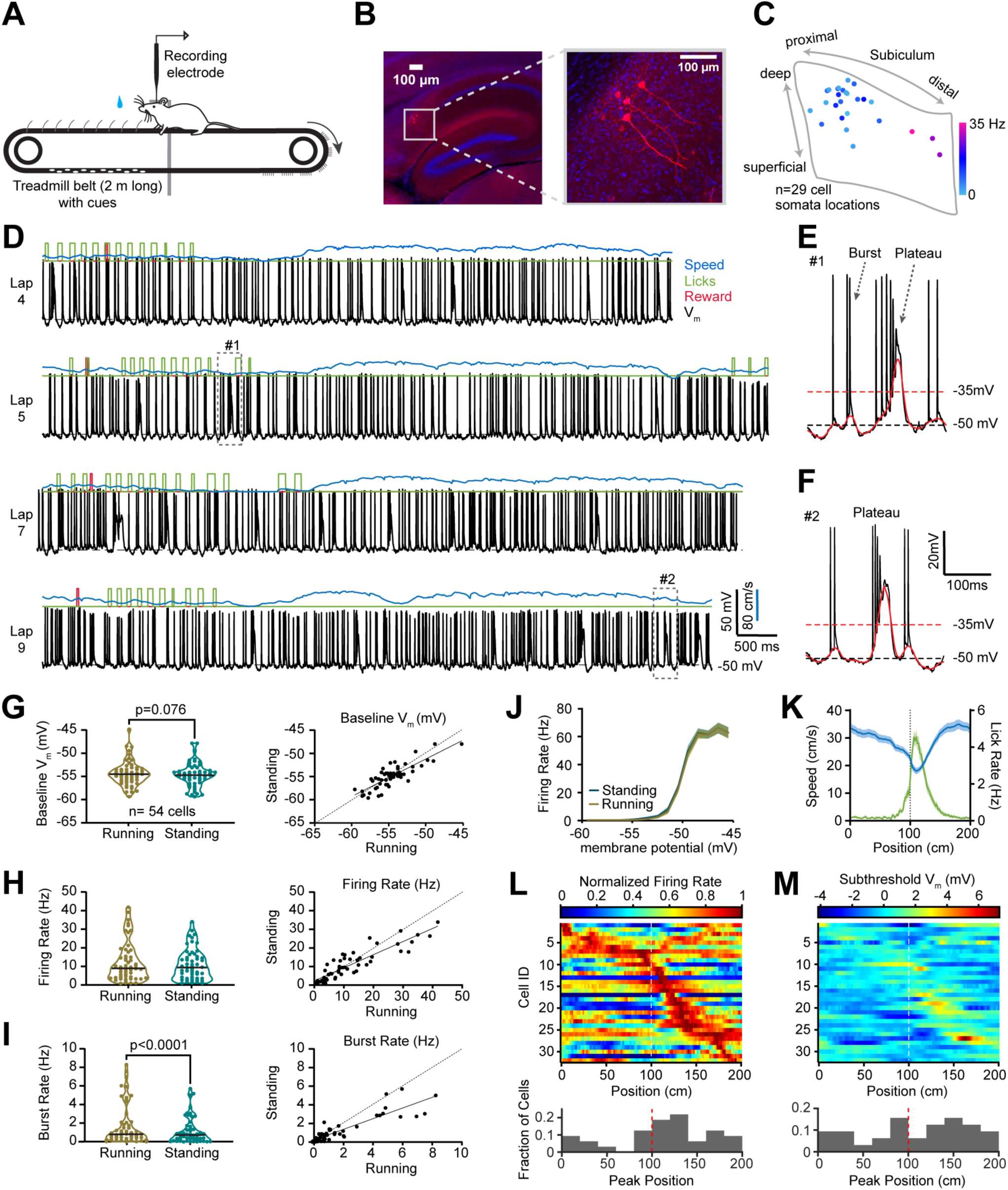
Whole-cell recordings of subiculum pyramidal neurons in behaving mice. (A) Schematic of the experimental configuration for whole-cell patch-clamp recordings in a head-fixed mouse running on a treadmill. (B) Left, biocytin-filled subiculum neurons (red) with DAPI (blue); gray box indicates the region magnified at right. (C) Dots indicate locations of recovered biocytin-filled neurons within subiculum boundaries (gray, sagittal); dot color denotes the mean firing rate across all the neurons recorded from the animal. (D) Example traces from a subiculum pyramidal neuron: raw membrane potential (V_m_) (black), running speed (blue), reward (red), and licks (green). “Lap” denotes one complete 2-m belt revolution. (E, F) V_m_ (black) traces corresponding to gray boxes #1 and #2 in panel D, showing single action potentials (APs), bursts, and plateaus. Plateaus are detected when the subthreshold V_m_ (red) crosses the −35 mV-threshold. (G-I) Baseline V_m_ (G), AP firing rate (H), and burst rate (I) during running vs. standing (Wilcoxon signed-rank; V_m_: p=0.076; firing rate: p=0.159; burst rate: p<0.0001; n=54). Left, violin plots (black line, median). Right, per-neuron scatter plots (standing vs. running). (J) Relationship between the V_m_ and AP firing rate during running vs. standing. (K) Running speed (blue) and lick rate (green) averaged over 2091 laps from 44 neurons. The dotted line indicates the location of the reward. (L-M) Top, peak normalized AP firing rate (L) and subthreshold V_m_ (M) heatmaps for individual neurons (n=32) sorted by peak firing position. Bottom, histograms of peak positions.

To assess whether subicular activity exhibited spatial structure in the fixed reward paradigm, we analyzed AP firing rate and subthreshold V_m_ as a function of position along the track (Fig. 1K–M). Individual neurons displayed heterogeneous spatial profiles, as expected from the literature^6,10,40,47–49^, with some cells showing peaks near the reward location and others firing broadly across the track. Approximately 59% of neurons (19 of 32) exhibited their peak firing rate within a ±40 cm window around the reward location (Fig. 1L). Subthreshold V_m_ showed weaker and less organized spatial modulation (Fig. 1M). Finally, to evaluate the contribution of movement-related variables to subicular activity in our paradigm, in which velocity and position are relatively correlated (Fig. 1K), we quantified the correlation between instantaneous running speed and AP firing rate and found that positively correlated, negatively correlated, and uncorrelated neurons were comparably represented (Supp. Fig. 1A–D). We also recorded from mice in a version of the paradigm in which the reward location was randomized on each lap (Supp. Fig. 1E-H). Neuronal firing rates were comparable between fixed and random reward conditions, whereas spatial information was reduced (Supp. Fig. 1E-F). A Generalized Linear Model (GLM) revealed reduced spatial coefficients with unchanged speed coefficients in the randomized condition (Supp. Fig. 1G–H). In summary, *in vivo* whole-cell recordings from dorsal subiculum neurons during navigation revealed frequent plateau firing alongside heterogeneous spatial and velocity modulation of subthreshold V_m_ and AP activity, consistent with the subicular network supporting a distributed neuronal representation^40^. These results also establish that our approach provides reliable intracellular access to identified subiculum neurons during active navigation.

### Subicular plateau occurrence is shaped by intrinsic and behavioral factors

To characterize the plateau events observed in subicular pyramidal neurons, we first formalized their detection based on the subthreshold V_m_ dynamics (Fig. 2A–C). After removing APs from the intracellularly recorded V_m_ and applying low-pass filtering, plateau events were identified as associated with sustained depolarizations that crossed −35 mV (Fig. 2A, Methods)^36^. Within individual neurons, the distribution of filtered V_m_ values showed a clear separation between laps with plateaus and those without, with divergence emerging around the −35 mV threshold (Fig. 2B). This separation was preserved when averaging across all recorded neurons (Fig. 2C), supporting a consistent operational definition of plateau events across the dataset. We next asked how regularly plateau events occurred and whether their incidence differed across behavioral states (Fig. 2D–G). We found that plateaus were observed in the majority of recorded neurons at baseline V_m_ (no current injection, 85%, 46/54) and occurred on about half the laps within individual neurons (Fig. 2D). Plateau rate (mean±SEM, running: 0.23±0.05 Hz; standing: 0.25±0.05 Hz) and the number of plateaus per 100 APs (mean±SEM, running: 1.5±0.25 Hz; standing: 1.8±0.28 Hz) were similar during running and standing periods (Fig. 2E). Plateau durations ranged from tens to hundreds of milliseconds and were significantly prolonged during standing compared to running (Fig. 2F, p<0.0001, Mann-Whitney U test). Finally, plateaus occurred across the full range of AP firing rates during running and standing (Fig. 2G), indicating that plateau generation is not restricted to times of overall elevated activity.

**Figure 2:**
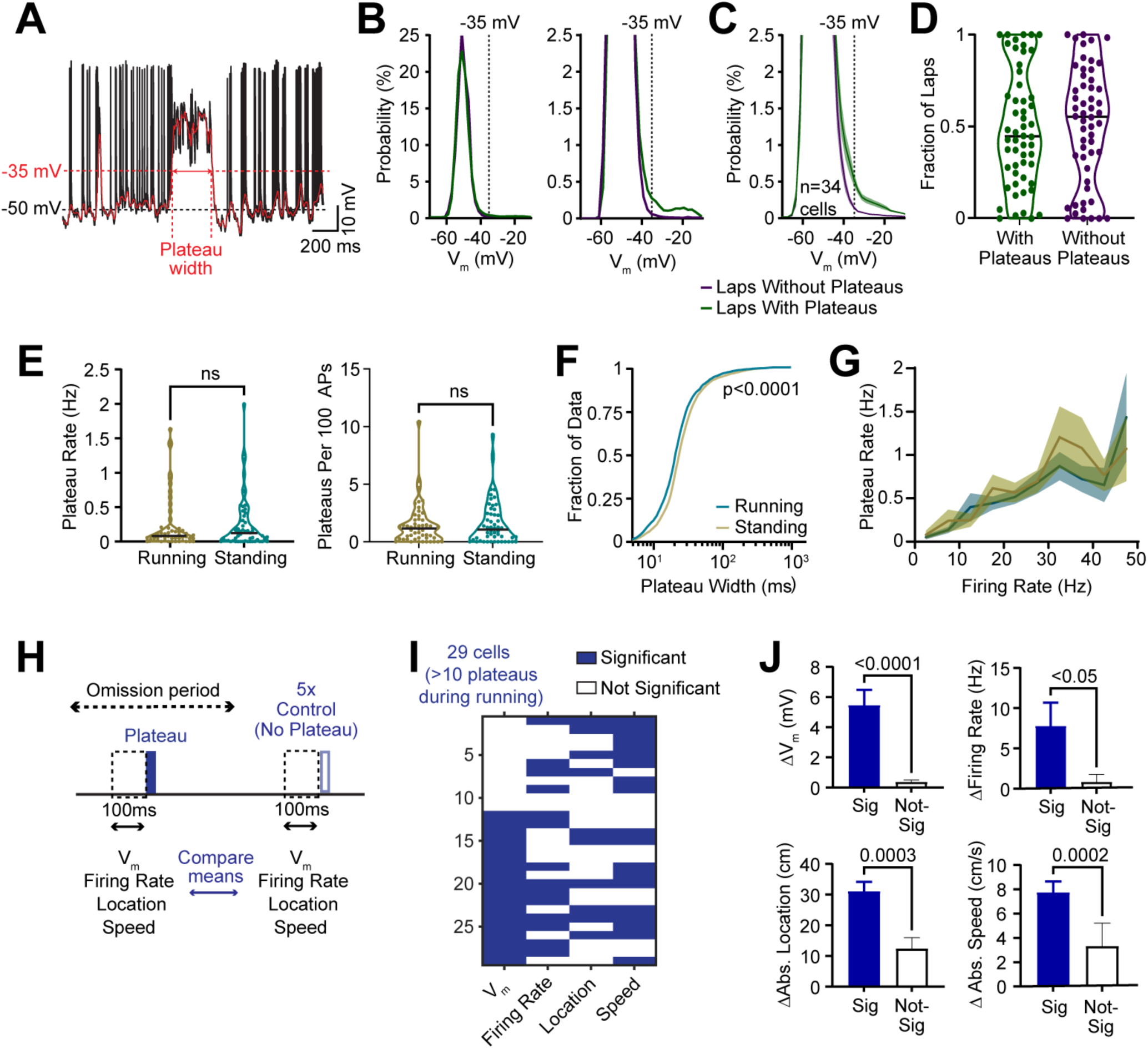
Characterization of subicular plateaus. (A) Raw V_m_ trace (black) from a subiculum pyramidal neuron illustrating plateau detection and width measurement. APs were removed and low-pass filtered (red), and plateau events were detected when crossing −35 mV. (B) Left, distribution of low-pass filtered V_m_ (red line in A) from an example neuron for running laps with (green) and without (purple) plateaus. Right, Same as left panel, but y-axis truncated to highlight the curve separation at −35 mV (dotted). (C) Same as B, averaged over all neurons with plateaus during running (n=34 neurons). (D) Fraction of laps with and without plateaus. (E) Mean plateau rate (left) and plateaus per 100 APs (right) during running vs. standing (Wilcoxon signed-rank test; n=54 neurons; plateau rate: p=0.905; plateaus per 100 APs: p=0.141). (F) Cumulative distributions of plateau widths during running vs. standing (Mann-Whitney U test, p<0.0001, running: n=3943 plateaus, standing: n=6910 plateaus). (G) Relationship between the plateau rate and the AP firing rate during running vs. standing. (H) Schematic of the univariate analysis. Mean V_m_, firing rate, location, and speed were computed in a 100 ms pre-plateau window (blue) and in five matched control windows outside a ±1 s exclusion period, then compared. (I) Significant (blue) and non-significant (white) differences between pre-plateau and control periods across variables (rank sum; p<0.05, n=29 cells with >10 plateaus during running). (J) Bar plots of mean differences between pre-plateau and control periods for cells with significant (blue) and non-significant effects; absolute differences were used for location and speed (Mann-Whitney U test). Bars, mean; error bars, SEM. The black line indicates the median in all violin plots.

Because behavioral variables are most consistently defined during locomotion, we restricted all subsequent analyses to plateaus occurring during running. To identify factors associated with plateau initiation under these conditions, we performed a univariate analysis comparing intrinsic and behavioral variables preceding plateau onset with matched non-plateau control periods (Fig. 2H). For each plateau, we quantified V_m_, firing rate, spatial location, and running speed within a 100 ms pre-event window, and compared these values with those obtained from randomly selected control time points from the same cells (Fig. 2H; Methods; Mann-Whitney U test). Across cells, the V_m_ preceding plateau onset emerged as the most consistent predictor of plateau occurrence (18/29 neurons), but neurons differed in which variables were predictive (Fig. 2I). For all parameters examined, cells exhibiting significant effects showed markedly larger differences between pre-plateau and control periods than cells without significant effects (Fig. 2J).

### Subicular plateaus exhibit voltage dependence and require NMDA receptor activation

To specifically assess how plateau firing depends on the V_m_, we measured plateau occurrence in n=13 neurons while biasing these neurons toward hyperpolarized (<-60 mV), intermediate (−60 mV to −50 mV), or depolarized (>-50 mV) V_m_ values via current injection through the patch pipette (Fig. 3A). Plateau events were readily observed at intermediate V_m_ values, increased in frequency during depolarization, and were strongly reduced during hyperpolarization (Fig. 3A-B). Thus, both the plateau rate and the number of plateaus per 100 APs per neuron increased systematically as cells were depolarized (Fig. 3B). In addition, a comparison of plateau widths across V_m_ conditions revealed that plateaus recorded during depolarized states tended to be significantly longer than those observed at intermediate V_m_ values (Fig. 3C). Together, these results indicate that plateaus are present at baseline or can be evoked by depolarization in all subicular pyramidal neurons, and that the V_m_ strongly regulates both the likelihood of plateau initiation and the duration of plateaus once triggered. The dependence of plateau occurrence on depolarization suggested a role for voltage-dependent conductances, such as NMDA receptors. We therefore tested whether plateau events require NMDA receptor activation by including the use-dependent NMDA receptor antagonist MK801 in the intracellular solution (Fig. 3D-H). MK801 significantly reduced the plateau rate and the number of plateaus per 100 APs (Fig. 3D-E). In addition, plateau durations were reduced in the presence of MK801, both when pooling plateau events across cells and when comparing median plateau width on a per-cell basis (Fig. 3F-G). Importantly, intracellular MK801 did not affect overall AP firing rate (Fig. 3H) or other neuronal properties, including baseline V_m_, burst rate, or number of APs per burst (Supp. Fig. 2). Together, these results demonstrate a critical role for NMDA receptors in plateau generation, suggesting a synaptic origin of subicular plateaus. These properties parallel the mechanisms underlying plateaus in CA1^33,34,36^, indicating shared cellular mechanisms with upstream hippocampal regions.

**Figure 3.**
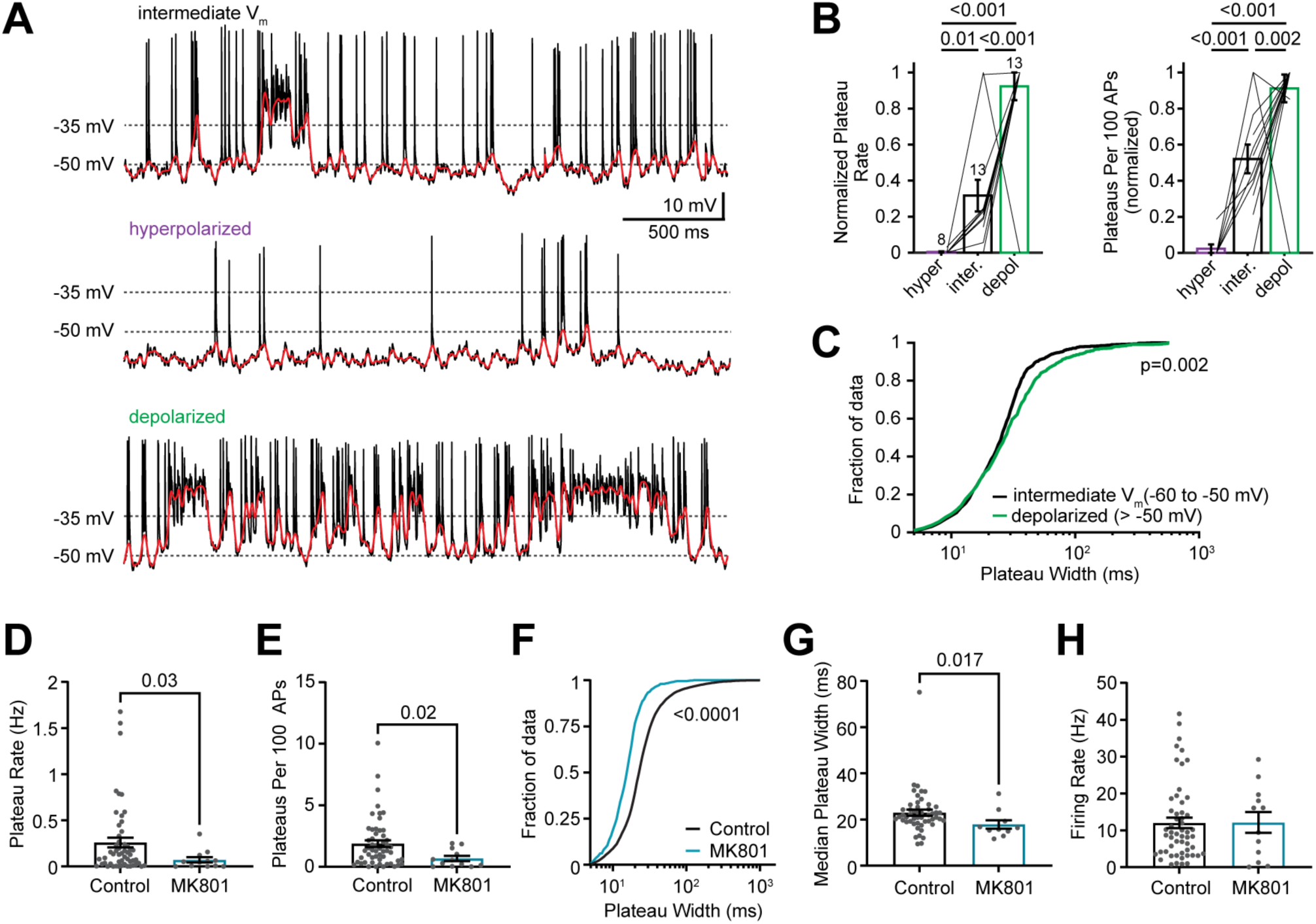
Subicular plateaus require NMDA receptor activation. (A) Representative pyramidal neuron V_m_ traces (black), with low-pass filtered V_m_ (red), for intermediate (−60 to −50 mV), hyperpolarized (<-60 mV), and depolarized (>-50 mV) conditions. (B) Normalized plateau rate (left) and plateaus per 100 APs (right) across V_m_ conditions (violet: hyperpolarized, black: intermediate, green: depolarized). Cell number per condition, and p-values (unpaired two-tailed *t*-test) are indicated; the same cells contribute to both measures. (C) Cumulative distribution of plateau widths for intermediate (n=877 plateaus, black) and depolarized (n=773 plateaus, green) V_m_ conditions (Mann-Whitney U test; p=0.002). (D-H) Data comparing control cells (black) and those with intracellular MK801 (teal) present; (D) plateau rate (p=0.03), (E) plateaus per 100 APs (p=0.02), (F) plateau width (p<0.0001), (G) median plateau width per cell (p=0.017), and (H) AP firing rate (p=0.908) (Mann Whitney U test, control: n=54, MK801: n=12). Data from running and standing periods were pooled for panels D-H. Data are shown as mean±SEM.

### Plateau activity clusters around the reward location

Next, we asked whether plateau events encode task-relevant information, such as the location of the reward. Therefore, we examined their spatial distribution relative to the fixed reward in 38 subicular neurons that exhibited at least 10 plateaus during running. To quantify task-relative organization while controlling for differences in event duration and dwell time (Supp. Fig. 3A), we measured “fraction plateau time” (the fraction of time the V_m_ exceeds the detection threshold of −35 mV, Supp. Fig. 3B, Methods) as a function of distance from the reward and from multiple control “anchor” positions (see Methods; Supp. Fig. 3C). Using this measure, analysis of all plateau events together revealed no significant reward-related clustering (Supp. Fig. 3D–E), indicating that any reward-related representation is likely restricted to a subset of plateau events.

Because synaptic inputs carrying distinct types of information impinge on subicular neurons at different distances from the soma and are, as a result, differentially attenuated on their way to the soma^50,51^, we reasoned that the somatic V_m_ preceding plateau onset could be used to discriminate plateaus associated with different task-relevant parameters. We tested this hypothesis by grouping plateau events based on the average subthreshold V_m_ in the 100 ms preceding onset, separating events occurring at relatively more hyperpolarized (“low V_m_”, −56 to −48 mV) versus more depolarized (“high V_m_”, −48 to −44 mV) values (Fig. 4A-F). An initial visualization of plateau locations suggested greater clustering of low V_m_ plateaus near the reward compared to high V_m_ plateaus (Fig. 4A–B). Importantly, all 38 neurons exhibited plateaus across the full range of pre-onset V_m_ values, with high V_m_ events comprising ∼42% and low V_m_ events comprising ∼58% of all plateaus (Fig. 4C-D). Plateau events in these two V_m_ ranges differed in several features. High V_m_ plateaus lasted on average longer (Fig. 4E) and were preceded by higher spike rates in the 1 s before onset (Fig. 4F). Furthermore, low V_m_ plateaus exhibited tighter phase locking relative to the extracellular theta oscillation (Fig. 4G), and a joint representation of plateau proportion as a function of this relative theta phase and distance from the reward confirmed the previously observed clustering of low V_m_ plateaus near the reward location (Fig. 4H).

**Figure 4:**
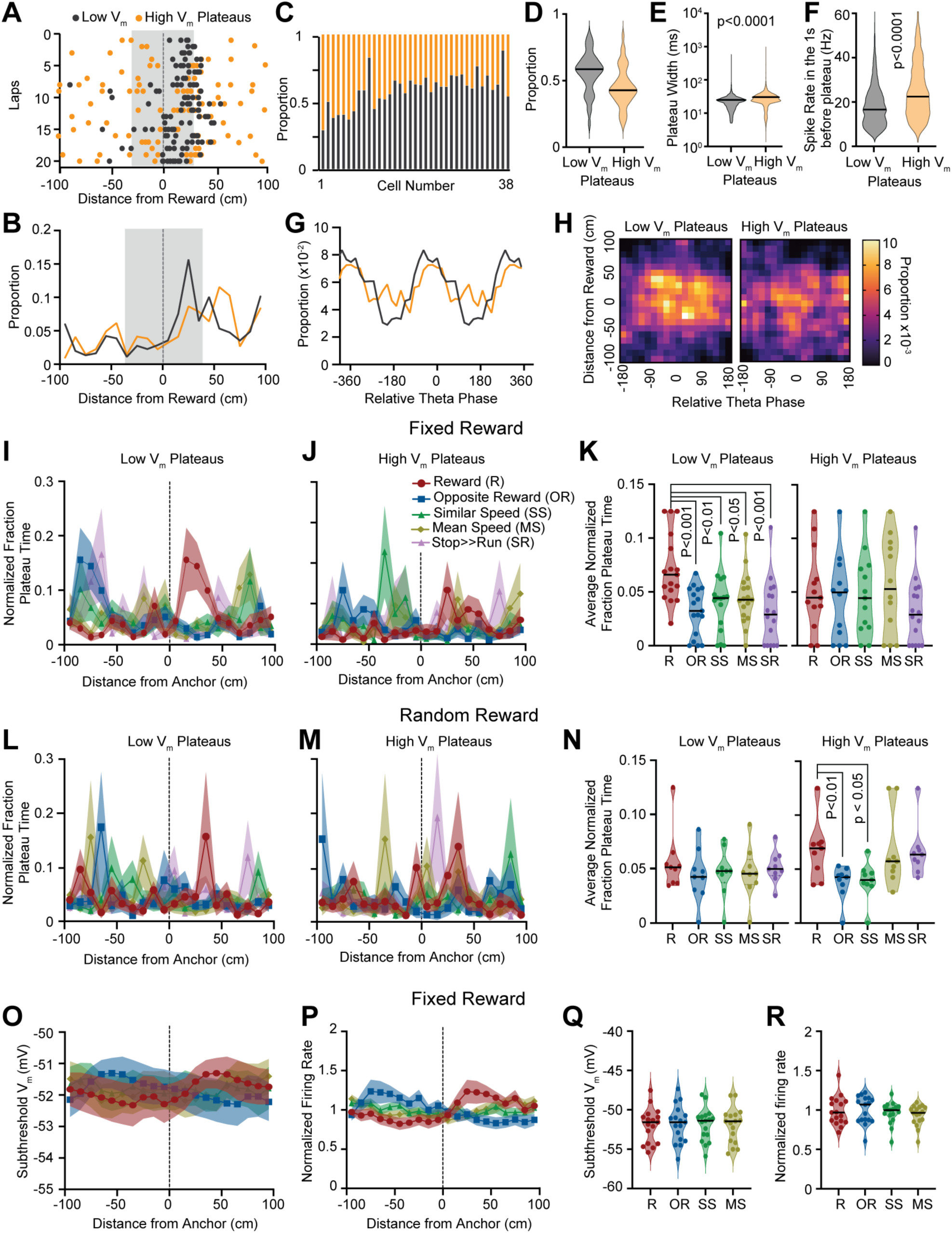
Subicular low V_m_ plateaus cluster around the reward location. Representative pyramidal neuron showing low (black) and high (orange) V_m_ plateau locations across laps and track position relative to reward. (B) Mean proportion of low (black) and high V_m_ (orange) plateaus across track position relative to reward (n=38 neurons); gray shading indicates the ±40 cm reward zone. (C) Proportion of low and high V_m_ plateaus in each of the 38 neurons. (D) Proportion of low and high V_m_ plateaus across all neurons. (E) Width of low and high V_m_ plateaus. (F) AP firing rate in the 1 second before the low and high V_m_ plateaus (Mann-Whitney U test). (G) Proportion of low and high V_m_ plateau onset as a function of LFP theta phase. The mean phase of the low V_m_ group is set to 0 degrees. (H) Heat maps of plateau proportions versus relative theta phase at onset (x-axis) and distance from reward (y-axis). Left, low V_m_ plateaus; right, high V_m_ plateaus. (I-K) Normalized plateau time in the fixed-reward condition. (I) Low V_m_ plateaus versus distance from reward (R, red circles) and movement-matched control anchors: opposite reward location (OR, blue squares), similar speed (SS, green triangles), mean speed (MS, brown diamonds), and stop→run transitions (SR, violet triangles). (J) same as (I) but for high V_m_ plateaus. (K) Quantification for (I) and (J). (L-N) similar to (I-K), but for the random reward condition. (O) and (P) Subthreshold V_m_ and normalized firing rate are plotted as distance from the 4 main anchors (R, OR, SS, and MS). (Q) and (R) Quantification of (O) and (P) respectively. The black line indicates the median in all violin plots. Detailed statistical information is listed in Supplementary Table 1.

To quantify this apparent spatial enrichment near the reward, we again used fraction plateau time and found that low V_m_ plateaus showed significantly higher values within a ±40 cm window around the reward location than within the same window around all control anchors (Fig. 4I, K). In contrast, high V_m_ plateaus did not show significant enrichment around the reward (Fig. 4J-K). To test whether this spatial organization depended on a stable reward location, we repeated the analysis in sessions in which reward delivery was randomized (Fig. 4L-N, n=9 neurons). Under these conditions, no clustering around the reward was observed when considering all plateaus (Supp. Fig. 3D–E), and neither low nor high V_m_ plateaus showed significant enrichment around the reward location relative to all the other control anchors (Fig. 4L-N), implying that the clustering depended on a reward that is expected. Finally, we asked whether the enrichment of plateau activity near the reward could be explained by changes in overall subthreshold V_m_ or AP firing rate (Fig. 4O-R). Neither subthreshold V_m_ nor normalized firing rate differed across the reward and movement-matched control anchors. Together, these analyses show that plateau activity is organized in a context-dependent manner: only plateaus preceded by relatively hyperpolarized V_m_ values accumulate near expected reward locations.

### Unsupervised clustering organizes plateaus by pre-plateau V_m_

To independently test whether pre V_m_ is a strong indicator of reward-aligned plateaus, we used an unsupervised clustering approach. We first performed a principal component analysis (PCA) that showed that pre-plateau V_m_ (pre V_m_), a Pearson’s correlation score against a global template (shape; see Methods), and either plateau duration or area (log-transformed) together explained more than 95% of the variance in the data (Supp. Fig. 4A-B). Because duration and area were highly correlated (r=0.91), we included only plateau area (log-transformed) in our analysis. Pairwise correlation analysis confirmed that pre V_m_, area, and shape were weakly correlated with one another (Supp. Fig. 4C), and both PCA and z-scored data indicated that k=2 clusters provided optimal separation (mean Silhouette score for PCA data: 0.62; mean Silhouette score for z-scored data: 0.564; Supp. Fig. 4D–E). Visualization using t-distributed stochastic neighbor embedding (t-SNE) shows two separable groups of plateau events (Fig. 5A). All neurons contributed plateau events to both clusters, although the relative proportion varied across cells (Fig. 5B, Supp. Fig. 4F). On average, cluster A comprised ∼60% of all plateaus, whereas cluster B contained ∼40%. Comparison of plateau features and morphology across clusters revealed that particularly pre V_m_ differed between clusters (Fig. 5C-G, Supp. Fig. 4H-I), which is also reflected by the highest Fisher’s linear discriminant ratio (Supp. Fig. 4G). We next examined whether the two plateau clusters differed in their spatial organization relative to the reward (Fig. 5H-K) and found that cluster A plateaus showed a pronounced enrichment around the reward location compared to control anchors (Fig. 5H, J). In contrast, Cluster B showed no statistically significant spatial modulation (Fig. 5I, K). Finally, we tested whether our pre-V_m_ grouping reflects extremes along a continuum, with the most hyperpolarized values biased toward reward. Using a sliding-window analysis of the pre-V_m_ distribution (20-percentile width, 10-percentile step), we found that plateaus preceded by more hyperpolarized V_m_ values showed greater reward selectivity, which progressively declined toward more depolarized V_m_ values (Fig. 5L). Thus, an unsupervised clustering approach converges with our earlier analyses (Fig. 4) in identifying pre-plateau V_m_ as a central organizing feature of reward-related plateau activity in the subiculum. We therefore returned to the same classification approach as used in Fig. 4 to determine how plateau activity in the subiculum adapts to changes in environmental features, such as the reward.

**Figure 5:**
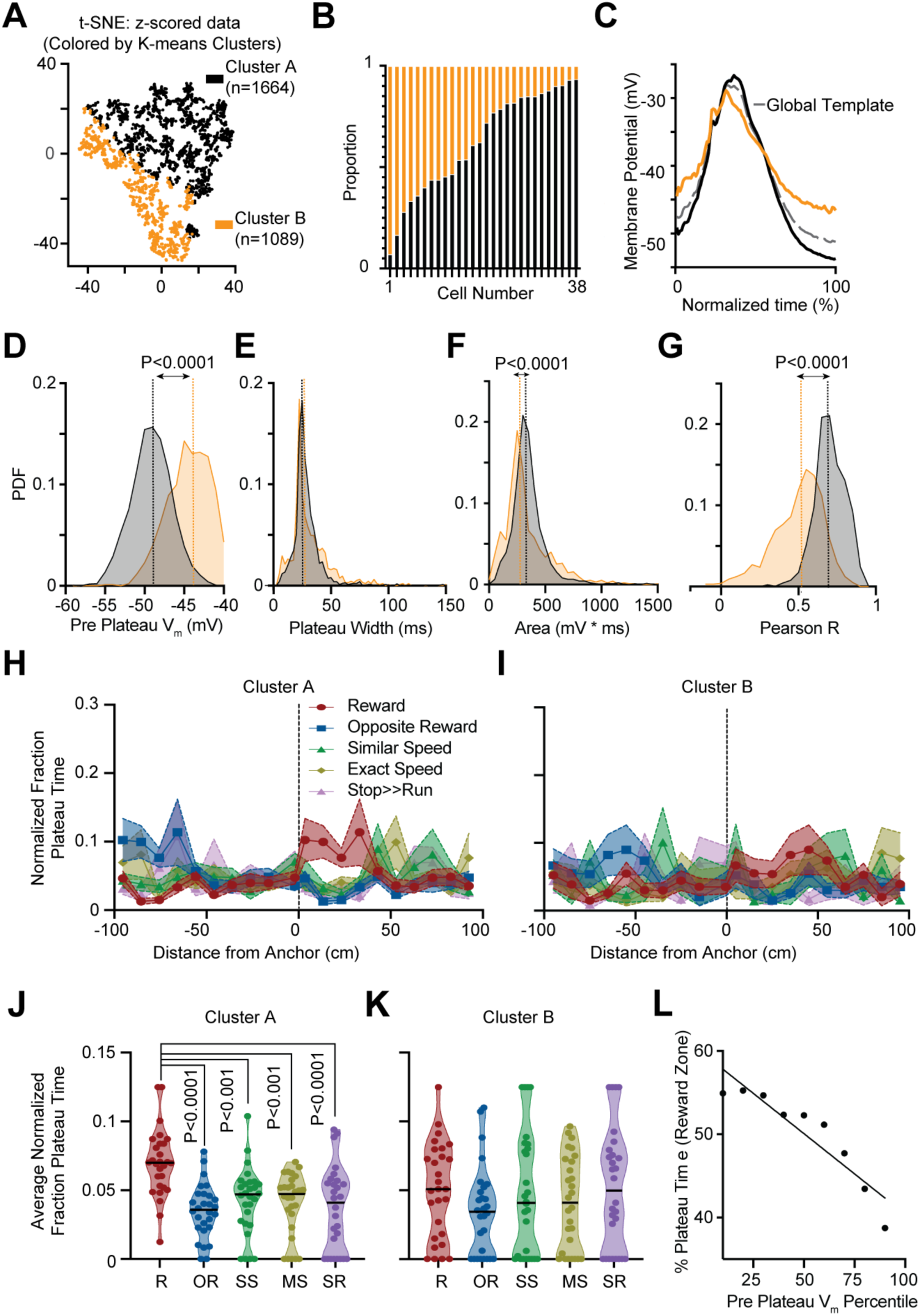
Unsupervised clustering identifies two classes of subicular plateaus. (A) t-SNE visualization of plateaus following k-means clustering based on pre-plateau V_m_ (Pre V_m_), plateau area (log-transformed), and plateau shape. Points are colored by cluster assignment (cluster A: black; cluster B: orange). (B) Proportion of plateaus assigned to cluster A and cluster B for each neuron. (C) Mean plateau waveforms for cluster A and cluster B, illustrating differences in pre-onset V_m_ (D-G) Probability Density Functions (PDFs) of pre-plateau V_m_ (D), plateau widths (E), area (F), and Pearson’s correlation coefficients (vs. global template) (G) for plateaus in cluster A and cluster B (Mann-Whitney U test). (H) Normalized fraction plateau time versus distance from reward (R, red circles) and movement-matched control anchors (opposite reward location: OR, blue squares; similar speed: SS, green triangles; mean speed: MS, brown diamonds; stop→run transitions: SR, violet triangles) for cluster A plateaus. (I) Same as (H), but for cluster B. (J, K) Quantification of normalized fraction plateau time within the ±40 cm reward zone for cluster A and cluster B plateaus. (L) Percentage plateau time in the reward zone as a function of pre-plateau V_m_ percentiles. Data are shown as mean±SEM unless otherwise indicated. The black line indicates the median in all violin plots. Detailed statistical information is provided in Supplementary Table 1.

### Plateau activity reorganizes after a change in reward location

To that end, we compared plateau occurrence before and after a 100 cm shift in reward location (half a lap; n=11 neurons). As illustrated by an example neuron, low V_m_ plateaus reorganized their spatial distribution following reward shift, whereas high V_m_ plateaus showed less reorganization (Fig. 6A). Using fraction plateau time across 11 neurons revealed that, before the shift, low V_m_ plateau activity was concentrated after the reward location (40 cm post-reward, Fig. 6B), as expected based on Figure 4. Following the shift, low V_m_ plateau activity increased around the new reward, with most low V_m_ plateaus occurring in the anticipatory region (40 cm pre-reward). High V_m_ plateaus, however, did not exhibit a comparable redistribution (Fig. 6C). To further quantify this finding, we defined spatial zones around the original and new reward locations (Fig. 6D). The “Learning Zone” was defined as the region around the new reward location after the shift, with the corresponding “Learning Control Zone” being the pre-shift reward location. Conversely, the “Memory Zone” was defined as the region surrounding the old reward location after the shift, with its control (“Memory Control Zone”) being the location that would later become the new reward site before the shift. Each zone was further subdivided into anticipatory and consumption periods. We first examined fraction plateau time during the consumption period and found that neither low nor high V_m_ plateau activity differed (Fig. 6E-G).

**Figure 6:**
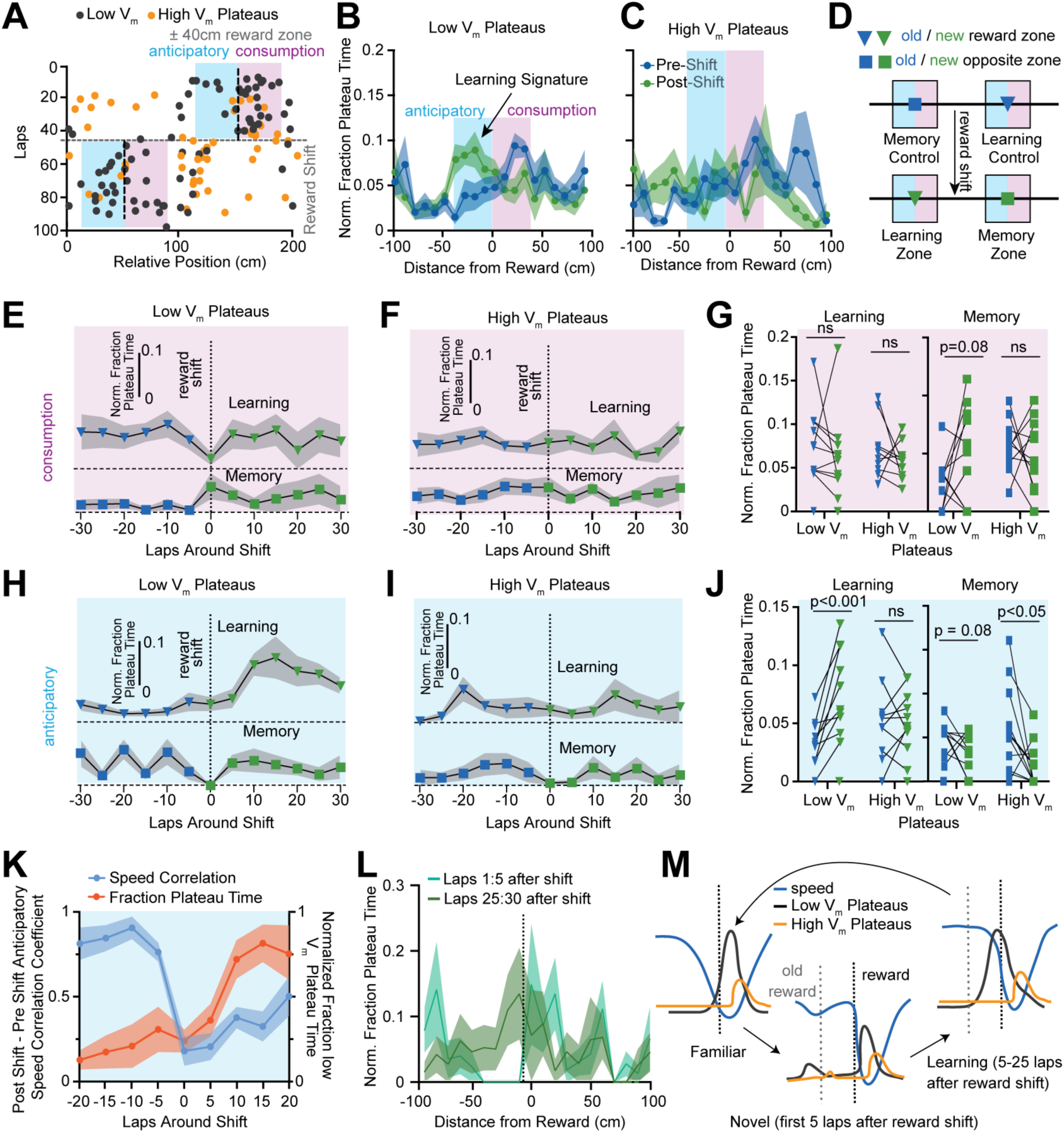
Subicular plateaus before and after reward location shift. (A) Plateau locations from a representative subicular pyramidal neuron with the reward location shifted after 45 laps. Low V_m_ plateaus (black), high V_m_ plateaus (orange), reward (vertical dashed line), reward shift (horizontal dashed line), anticipatory region (blue), and consumption region (pink). (B) Distribution of normalized fraction plateau time for low V_m_ plateaus across all 11 neurons before and after reward shift. Familiar location before reward shift (blue) and novel location after reward shift (green). (C) same as (B), but for high V_m_ plateaus. (D) Schematic illustrating the learning zone (green triangle), the learning control zone (blue triangle), the memory zone (green square), and the memory control zone (blue square). Each is split into equal-sized anticipatory (blue) and consumption (pink) regions. (E) Change in normalized low V_m_ plateau time in the consumption region from learning control to learning zone and from memory control to memory zone (5-lap averages); learning control/zone baseline is offset for clarity. (F) same as (E) but for high V_m_ plateaus. (G) Quantification of (E) and (F). (H) and (I) same as (E) and (F), but for the anticipatory regions. (J) Quantification of (H) and (I). (K) Relationship between anticipatory low V_m_ plateau activity and running behavior around the reward shift. Blue, mean speed correlation (5-lap bins vs. pre-shift average; left axis). Orange, normalized low V_m_ plateau time in the anticipatory region (5-lap bins; right axis). (L) Change in low V_m_ plateau fraction during the first 5 laps after reward shift (cyan) versus laps 25-30 post-shift (green). (M) Summary schematic of plateau dynamics in familiar and reward-shift conditions. All line plots are plotted as mean±SEM, unless noted otherwise. Detailed statistical information is listed in Table 1.

We next focused on the anticipatory period preceding reward delivery (Fig. 6H-J). During this period, low V_m_ plateaus showed a significant increase in fraction plateau time within the Learning Zone compared to the Learning Control Zone following the reward shift (Fig. 6H, J). This increase emerged with a delay, peaking approximately 10-15 laps after the shift. No corresponding change was observed in the anticipatory Memory Zone. High V_m_ plateaus, however, did not show increased activity in the anticipatory Learning Zone; instead, high V_m_ fraction plateau decreased significantly in the Memory Zone following the shift (Fig. 6I-J). Together, these results demonstrate a reorganization of anticipatory low and high V_m_ plateau activity associated with learning a new reward location. To rule out that this change in plateau activity reflected residual spatial memory of the former reward location, we compared the fraction plateau time at the original reward site with the same physical location after the reward shift, separately for low and high V_m_ plateaus, and found no differences in either the anticipatory or consumption regions (Supp. Fig. 5).

Finally, we asked whether the delayed increase in low V_m_ plateau activity in the anticipatory Learning Zone, i.e., the most prominent reorganization observed, was associated with behavioral adaptation. We correlated the running speed profile in the 100 cm preceding the new reward location after the shift with the speed profile in the 100 cm preceding the old reward location before the shift. Immediately after the reward shift, this correlation was low but gradually increased across laps as animals adjusted their behavior to the new reward location (Fig. 6K). Notably, this behavioral change occurred in parallel with the delayed increase in fraction plateau time of low V_m_ plateaus in the anticipatory Learning Zone. Consistent with this relationship, when post-shift laps were divided into early (laps 1-5) and late (laps 25-30) periods, the spatial peak of low V_m_ plateau activity shifted from just after reward delivery in early laps to approximately 20 cm before reward delivery in late laps (Fig. 6L). This progression was specific to low V_m_ plateaus, as high V_m_ plateaus did not exhibit a comparable shift across laps (Fig. 6M, Supp. Fig. 5C).

## Discussion

The subiculum occupies a unique position as one of the major outputs of the dorsal hippocampal formation, yet its single-cell operating principles during behavior have remained poorly defined. To our knowledge, this study provides the first systematic *in vivo* whole-cell characterization of subicular pyramidal neuron activity during goal-directed navigation. In line with previous reports^14,40^, we show that subicular neurons exhibit unusually high firing rates and a pronounced tendency to burst. Our intracellular recordings reveal further that this burst firing is frequently driven by sustained, NMDA-dependent depolarizations, establishing plateau activity as a prominent cellular mechanism of subicular output during behavior. Because plateau events are defined by large, sustained subthreshold depolarizations underlying burst firing, they cannot be distinguished from other forms of burst activity using extracellular methods. Thus, intracellular recordings are essential for resolving the mechanistic diversity of subicular bursting during behavior.

Plateaus were observed during both running and standing, implying that subicular plateau activity reflects internal network dynamics or global brain states that shape hippocampal output beyond moment-to-moment locomotor behavior. Unlike subicular bursts described *in vitro*, which are primarily driven by voltage-dependent conductances and do not require synaptic input^20,21,25,27^, the plateaus observed here depend on NMDA receptor activation, indicating a strong synaptic contribution. In addition, whereas CA1 plateaus are typically spatially sparse and tightly linked to place field firing^30,35,36,52^, subicular plateau events were more abundant and broadly distributed across space, suggesting that the subiculum in behaving mice operates in a regime that uses reliable, burst-focused signal transmission, consistent with its role as a major hippocampal output region. Consistently, plateau-associated AP bursts are well positioned to generate strong, temporally structured output signals, and the plateau-related depolarizations may influence transmission more subtly, as subthreshold membrane potentials have been shown to propagate along central axons and modulate AP-evoked release^53–55^. Thus, subicular neurons may use a combined signaling mode, in which burst firing provides discrete, temporally structured signals while the underlying depolarized V_m_ dynamically modulates spike-triggered transmission.

Our most striking finding is that the V_m_ preceding plateau onset reveals systematic differences in the behavioral associations of plateaus. Notably, these differences emerged when considering all neurons, regardless of their feature selectivity, suggesting that this feature is widespread among subicular neurons. Plateau events occurring at relatively more hyperpolarized V_m_ values preferentially clustered near reward locations and were selectively recruited during learning of a new reward position, whereas plateau events occurring at more depolarized V_m_ values were more broadly distributed. To assess whether this organization could be recovered without imposing V_m_-based criteria, we applied an unsupervised clustering approach to plateau features (Fig. 5). This analysis converged with our V_m_-based grouping (Figs. 4, 6), and together, these results implicate intracellular V_m_ state as a crucial organizing variable for subicular plateau output.

Notably, the distinction between low and high V_m_ plateaus should not be taken as evidence for discrete plateau types, although we cannot fully exclude that possibility. Instead, our data support a continuum of cellular or network activation states in which individual neurons may dynamically bias their signaling toward different behaviorally relevant features, such as reward-related vs. non-reward information. Differences in duration, firing context, and theta phase locking between low and high V_m_ plateaus may affect how downstream circuits respond to subicular activity^56^, without requiring discrete plateau classes. Given the broad projection targets of the subiculum^5,6^, this output modulation could provide a mechanism by which reward-related signals are emphasized under specific internal activation states. In this context, it will be particularly interesting to relate subicular plateau activity to cell types previously identified in downstream regions, such as goal representations described in entorhinal and related cortices^57,58^, which may be especially sensitive to the temporal structure of hippocampal output. Together, our findings establish plateau activity as a central mechanism structuring hippocampal output at the level of the subiculum. At the same time, this work raises important questions about the cellular and circuit mechanisms that generate these events, how downstream regions interpret subicular output to guide behavior, and whether similar state-dependent plateau-based signaling modes operate in other brain circuits that exhibit plateau events.

## Acknowledgements

We thank Francisco Mello for technical assistance. We thank Eve Marder and Gina Turrigiano for valuable discussions, and Yangqi (James) Jiang and Natalie MacKinnon-Booth for help with animal training and histology in the early phases of this project. This work was supported by the NIH 1DP2 MH136393 (CG), NIH T32 NS007292 (MA), the Smith Foundation (CG), and the Swartz Foundation (AB).

## Author Contributions

AB, MA, and CG designed the research. AB and MA performed *in vivo* recordings. AB, MA, and CG analyzed the data and wrote the manuscript.

## Methods

All experiments were performed according to methods approved by the Institutional Animal Care and Use Committees at Brandeis University (protocol #22022 and #25001).

### Surgery and training

All experiments were performed in male and female mice aged 5-10 weeks by an experimenter who was not blind to the experimental condition. Animals were housed on a reversed 12 h light/12 h dark cycle. All surgical procedures were conducted under deep isoflurane anesthesia. After local application of a topical anesthetic and betadine, the scalp was removed, and the skull was exposed, cleaned, and leveled. Stereotaxic coordinates for future craniotomies were marked at 3.0 mm caudal and 2.25 mm lateral for whole-cell recordings, and at 3.5 mm caudal and 2.75 mm lateral for local field potential (LFP) recordings. The skull was then coated with cyanoacrylate glue (Loctite), and a custom-made titanium head bar with an opening over the hippocampus was then attached to the skull using dental acrylic (Ortho-Jet, Lang Dental). Five to seven days after head-bar implantation, mice were placed on water restriction. Following at least 3 days of water restriction, animals underwent handling and experimenter familiarization for at least 5 days, followed by the training to run on a 2 m linear treadmill for 6-12 days. Training was conducted during the animals’ dark cycle. The self-propelled velvet treadmill belt (McMaster-Carr) was equipped with visual and tactile cues, dividing it into three distinct sections^36,59^. During training, mice received a 10% sucrose/water reward via a 3D-printed lick port. Photoelectric sensors in the lick port and beneath the treadmill detected licks and lap completion, respectively, while an encoder (Broadcom) attached to the treadmill axle measured running speed. All sensors, the solenoid valve (The Lee Company), and the encoder were controlled using Bpod (Sanworks) and custom MATLAB software. Most animals were trained under a fixed-reward paradigm on the cued belt, with reward delivery at a constant belt location. In a subset of mice, the reward location was shifted by 100 cm (half a lap) after an initial set of laps during a recording. Another subset was trained in a random reward paradigm (belt with cues), receiving one reward per lap at a pseudo-randomly selected location. Animals were considered ready for electrophysiological recordings once they consistently completed more than 100 laps within 60 min or exhibited stable daily performance across two to three consecutive days. On the day before electrophysiological recordings, animals were anesthetized with either isoflurane or ketamine/xylazine, and two small craniotomies (∼0.75 mm in diameter), i.e., one for the patch electrode and one for the LFP electrode, were made at the previously marked locations, leaving the dura intact. Craniotomies were sealed with silicone elastomer (Kwik-Cast, World Precision Instruments), and recordings were performed over the subsequent 1-2 days.

### *In vivo* electrophysiology

For each animal, the approximate depth of the subicular pyramidal cell layer was identified using LFP recordings. For this, glass electrodes (1.5-3 MΩ) were filled with 0.9% NaCl and mounted vertically on a micromanipulator (Luigs & Neumann), and the LFP signal was monitored using an audio amplifier as the electrode was advanced through the cortex. As previously described^36,52^, the pyramidal cell layer was identified by the presence of theta-modulated AP activity, typically at depths of 1100–1400 μm below the brain surface. This electrode was then removed to make space for the whole-cell patch electrodes (see below). For simultaneous LFP recording during whole-cell patch-clamp experiments, a second extracellular electrode was then mounted on a separate micromanipulator (Narishige) at a 60° angle relative to bregma and advanced through a second craniotomy until the pyramidal cell layer was detected. Whole-cell patch electrodes (7-12 MΩ) were filled with an intracellular solution containing (in mM): 134 potassium gluconate, 6 KCl, 10 HEPES, 4 NaCl, 0.3 Mg-GTP, 4 Mg-ATP, and 14 Tris-phosphocreatine; biocytin 2% was included in all recordings. In a subset of neurons (n=12), the use-dependent NMDA receptor antagonist MK-801 (1-2 mM) was added to the intracellular solution. Patch electrodes were advanced into the subiculum under positive pressure (7-9 psi), which was reduced to 0.25-0.35 psi upon entry into the target region. Cells were identified by reproducible increases in electrode resistance. To examine the relationship between V_m_ and plateau activity, sustained current was injected through the recording pipette in a subset of neurons (n=13) to adjust V_m_ toward hyperpolarized (−70 to −60 mV), intermediate (−60 to −50 mV), or depolarized (−50 to −35 mV) ranges. All recordings were performed in current-clamp mode using an AM Systems Model 2400 patch-clamp amplifier, and LFP signals were acquired with an AM Systems Model 1700 differential amplifier. Signals were digitized at 20 kHz using a NIDAQ-based acquisition system (National Instruments, PCIe-6341) and acquired with MATLAB-based Wavesurfer software (version 0.982, Janelia). Interneurons were identified based on spike waveform characteristics and subthreshold V_m_ dynamics and were excluded from all analyses.

### Histology

To enable anatomical identification of recorded neurons, 2% biocytin was included in the intracellular recording solution. Following electrophysiological experiments, animals were transcardially perfused with 4% paraformaldehyde (PFA). Sagittal brain sections were then prepared and processed with Alexa Fluor 594–conjugated streptavidin to visualize biocytin-filled neurons and confirm their location within the subiculum.

### Data analysis

Intracellularly recorded V_m_ and simultaneous behavioral data were acquired at 20 kHz. Action potentials were detected based on the maximum upstroke velocity: a spike peak was identified when the first derivative of the voltage trace (dV/dt) exceeded a fixed threshold of 20 V/s (corresponding to 0.001 V/sample at 20 kHz), with a minimum inter-peak interval of 30 samples (1.5 ms). To correct for baseline drift, V_m_ traces were adjusted on a lap-by-lap basis. For each spike, the voltage threshold was first determined. The average deviation of the most hyperpolarized 5% of these thresholds from a target value of −50 mV was then calculated and subtracted from the entire V_m_ trace for that lap. Subthreshold membrane voltage was obtained by removing AP-associated voltage transients and interpolating across the resulting gaps. Specifically, for each detected spike, the voltage trace from 0.25 ms before to 6.4 ms after the spike peak was removed and replaced by linear interpolation using the subsequent available data point. The baseline subthreshold V_m_ for each neuron was defined as the mode of the resulting subthreshold voltage distribution, computed using 0.2 mV bins. Bursts were defined as events in which at least two APs occurred within an interspike interval of less than 10 ms. Plateaus were detected using a threshold-crossing criterion applied to the subthreshold V_m_ trace. Plateau onset and offset were defined as the time points at which a 401-point moving average of the subthreshold V_m_ crossed a −35 mV threshold. Only events longer than 5 ms were included, and only neurons completing ten or more laps were included in the analysis. Running periods were defined as epochs during which the animal’s running speed exceeded 4 cm/s. Unless otherwise noted, all analyses were restricted to data acquired during running.

### Quantification of plateau activity as a function of Vm

Continuous recording segments of at least 10 s duration were identified, including periods at baseline V_m_ (no current injection) as well as periods during which V_m_ was adjusted by sustained current injection. For each segment, V_m_ was quantified as the mode of the V_m_ distribution, computed from a histogram with a bin size of 0.2 mV. Within each segment, the number of plateau events and APs was quantified using the detection criteria described above. Plateau activity was characterized by its absolute event frequency and the number of plateaus per 100 APs. To enable comparisons across neurons with differing firing rates, both measures were normalized to the maximum value observed within each neuron before population-level analysis.

### Spatial Information and speed tuning analysis

Spatial information was calculated as previously described^60^. To assess the statistical significance of the relationship between running speed and instantaneous firing rate, we used a circular time-shifting procedure^61^. This approach tests the null hypothesis of temporal independence by generating a null distribution of correlation coefficients from 2,000 shuffled datasets. Specifically, the firing rate time series was circularly shifted by a random offset (30–100 s) and correlated with the unshifted running speed vector. The observed correlation was considered significant if it fell outside the central 98% of the null distribution (i.e., below the 1st percentile for negative correlations or above the 99th percentile for positive correlations).

### Univariate analysis

To understand the conditions that predicted the occurrence of plateaus, we performed a univariate analysis for each cell individually. To prepare the dataset for univariate analysis, we first aggregated event data from cells that exhibited more than ten plateaus during the running period. For both plateaus and non-plateau (control) events, specific features were calculated within a fixed 100 ms pre-event window (120 to 20 ms before event onset). Non-plateau events were randomly sampled from running periods, ensuring a 1s buffer from any actual plateau and maintaining a 5:1 non-plateau-to-plateau ratio whenever possible for balanced sampling. The dataset’s feature means within the 100 ms window were then used to compare the distributions of individual features between the “plateau” and “non-plateau” groups, cell by cell, using the Mann-Whitney U test. Cells recorded during the random reward condition and the laps after reward shift were excluded from this analysis.

### Generalized Linear Model (GLM)

To determine the relationship between instantaneous speed and firing rate, we performed a cell-by-cell Generalized Linear Model (GLM) analysis using a Poisson distribution and a canonical link function. The instantaneous firing rate served as the response variable, modeled as a function of the predictors, speed, and place (position on the track). Before fitting, all continuous predictor variables were rescaled to standardize their range and ensure the coefficients were directly comparable. For each fitted model, the resulting coefficients were extracted for comparison.

### Low V_m_ and high V_m_ plateaus

After plateau detection and subthreshold V_m_ extraction (as described above), we analyzed the subthreshold V_m_ during the 100 ms preceding each plateau event. 95% of all plateaus had a subthreshold V_m_ range between −44 mV and −56 mV. In a few instances, two plateaus occurred right after each other such that the 100 ms before the second plateau included all or some of the width of the first plateau. In this case, only the first plateau was included as it was not possible to determine the subthreshold V_m_ preceding it. All analyzed plateaus were then split into two groups based on their subthreshold V_m_: the highest ∼33% of the range (−44 mV to −48 mV) were categorized as high V_m_ plateaus, while the lowest ∼67% of the subthreshold V_m_ range (−48 mV to −56 mV) were categorized as low V_m_ plateaus. More than 88% of low V_m_ plateaus had a subthreshold V_m_ of −48 mV to −52 mV (the higher 50% of their range).

### Plateaus around reward and movement anchors

To assess whether plateau events cluster around the reward location, we compared reward-centered plateau distributions to multiple control anchor locations that should not exhibit similar clustering if plateaus are genuinely reward-related. These control anchors included: (1) the location opposite the reward (Opposite Reward; OR); (2) the 0.5 s window within the same lap with maximal speed correlation to the 0.5 s window surrounding the reward (Similar Speed; SS; only laps with a maximum correlation coefficient > 0.5 were included); (3) the 0.5 s window with mean running speed most similar to that around the reward (Mean Speed; MS; only laps in which the detected anchor differed by <5% from the reward-associated mean speed were included); and (4) the point at which the animal resumed running after stopping (Stop→Run; SR). In some recordings, the SR control was omitted if insufficient laps contained Stop→Run events located more than 50 cm from the reward. To avoid confounds related to proximity to the reward itself, all control anchors were constrained to be at least 50 cm away from the reward location on each lap. Anchor locations were computed independently for each lap, with a maximum of one anchor per lap for each control condition. For each anchor, the number of plateaus occurring within 10 cm spatial bins centered on the anchor was counted and divided by the number of laps with successful anchor detection, yielding the average number of plateaus per bin (“bin proportion”). Bin proportions were then normalized by dividing by their sum to obtain the normalized plateau probability for each spatial bin. This normalization allowed pooling of data across neurons. To control for differences in dwell time across spatial locations, we additionally computed the “Fraction Plateau Time” as the total time spent at V_m_ values >-35mV in plateau events within a bin divided by the total dwell time in that bin (Supp. Fig. 3B). This measure was similarly normalized by dividing the summed Fraction Plateau Time within the target bins by the total Fraction Plateau Time across all bins, enabling comparisons across recordings.

### Analysis of plateau activity associated with reward location shift

This analysis was performed using the same approach described above, but restricted to the old and new reward anchors, as well as to the locations where the new reward would have been delivered and where the old reward had previously been delivered. For this analysis, the reward-centered region (target bins) was further subdivided into anticipatory (40 cm preceding the anchor) and consumption (40 cm following the anchor) periods. To assess changes in plateau clustering during new learning, the original reward zone was used as a speed-matched control for the new reward zone. Similarly, the location opposite the old reward was used as a speed-matched control for the location opposite the new reward to test for memory-related effects on plateau clustering at the former reward location. To quantify the progression of plateau clustering across the session, laps were grouped into consecutive blocks of five centered on the reward-shift lap, and the analyses described above were applied to each block.

### Theta alignment

To determine the theta phase associated with plateau initiation (Fig. 4G-H), we computed the relative instantaneous phase of LFP theta oscillations at the onset of each detected plateau. First, the LFP signal was restricted to running periods and downsampled from the original acquisition rate (20 kHz) to 200 Hz. The down-sampled signal was bandpass-filtered in the theta range (4–8 Hz) using a zero-phase digital filter, then resampled to the original sampling rate to preserve precise alignment with plateau onset times. The analytic representation of the theta-filtered signal was obtained using the Hilbert transform, and instantaneous phase values (−π to π) were extracted for all plateaus. To combine data across neurons, circular phase distributions from individual recordings were pooled. For normalization, the mean phase of low V_m_ plateaus for each cell was defined as the reference (0 degrees), and phases for both low and high V_m_ plateaus were calculated relative to this reference. Linear histograms were then constructed to display the normalized proportion of plateaus as a function of this relative theta phase using 20° bins.

### Calculation of Area and Shape

For each plateau, the V_m_ trace between the −35 mV threshold crossings, including a 25 ms flanking window before and after the event, was extracted. All traces were then spike-filtered to isolate the low-frequency waveform. For area measurements, only the V_m_ segments between the two −35 mV threshold crossings were analyzed. After subtracting −35 mV from each point, the remaining trace was integrated, and the area was calculated as mV*ms. For the shape analysis, the full spike-filtered trace, including the 25-ms flanking regions, was used. The duration and amplitude of each event were normalized to a 0–100% scale, and the pointwise mean of these normalized waveforms was defined as the global template. A shape score for each plateau was then calculated as the Pearson correlation coefficient (r) between the individual event and the global template. This approach emphasizes morphological similarity independent of absolute amplitude or duration.

### Principal component analysis

To reduce dimensionality and mitigate multicollinearity, a Principal Component Analysis (PCA) was performed on the extracted variables (Pre V_m_, Area, Duration, and Shape score). Prior to the PCA, variables with heavy-tailed distributions (duration and area) were log-transformed (log10), and all variables were standardized (z-scored) to ensure equal weighting. The minimum number of principal components required to explain at least 95% of the cumulative variance was retained. The component scores from this reduced subspace were used as the input for the subsequent clustering analyses.

### K-means clustering and t-SNE visualization

The uncorrelated features (pre V_m_, log-area, and shape) were z-scored prior to clustering. To determine the optimal number of clusters (k), we evaluated the sum of squared errors and silhouette scores for k=1-6. Based on these diagnostics, we selected k=2. Clustering was performed using the k-means algorithm (squared Euclidean distance) with 10 replicates to minimize initialization bias. To visualize the structure of the event feature space in two dimensions, we applied t-distributed stochastic neighbor embedding (t-SNE) to the standardized feature matrix. t-SNE was used solely for visualization to assess the separation of clusters identified by k-means and did not influence the clustering itself.

