## Supplemental Information for "Subicular Plateaus Signal Reward Locations during Goal-Directed Behavior"

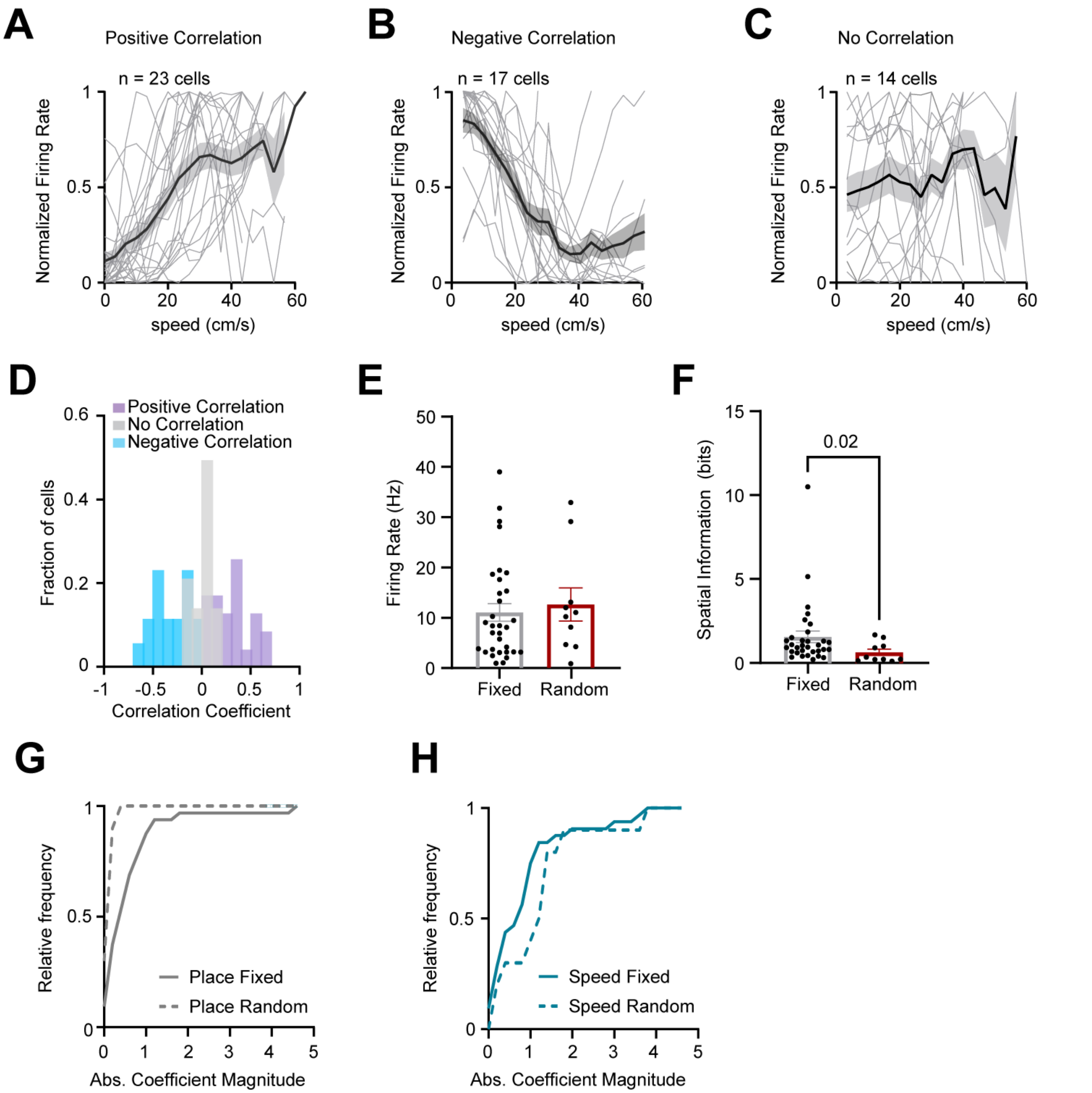


**Supplemental Figure 1: Characterization of subicular pyramidal neurons during goal-directed fixed and random reward conditions.** (A-C) Speed-firing rate tuning curves for neurons with positive (A), negative (B), or no correlation (C). Individual neurons (light gray), mean±SEM (dark gray). (D) Histogram of Pearson correlation coefficient between instantaneous running speed and AP firing rate for neurons with positive, negative, and no correlation. (E) Comparison of AP firing rate between fixed and random reward conditions (Mann-Whitney U test, p=0.512, fixed reward: n=32 neurons, random reward: n=10 neurons). (F) Same as E, but for spatial information (Mann-Whitney U test, p=0.024, fixed reward: n=32 neurons, random reward: n=10 neurons). (G) Cumulative distribution of GLM place coefficients for fixed (solid line) and random reward (dotted line) conditions. (H) Same as G, but for speed coefficients.

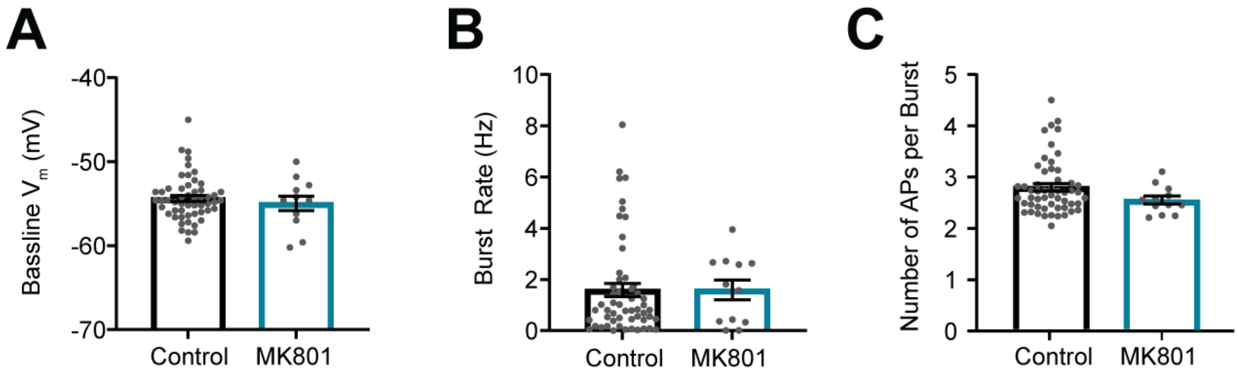


**Supplemental Figure 2: Baseline $V_m$ and firing properties during MK801 (NMDA receptor antagonist) application.** (A-C) Data comparing control pyramidal neurons and neurons recorded with intracellular MK801 present for (A) baseline $V_m$ ($p=0.946$), (B) burst rate ($p=0.773$), (C) number of APs per burst ($p = 0.125$) (Mann-Whitney U test, n=54 control, n=12 MK801 cells). Data from running and standing periods were pooled for panels A-C. Data are shown as mean±SEM.

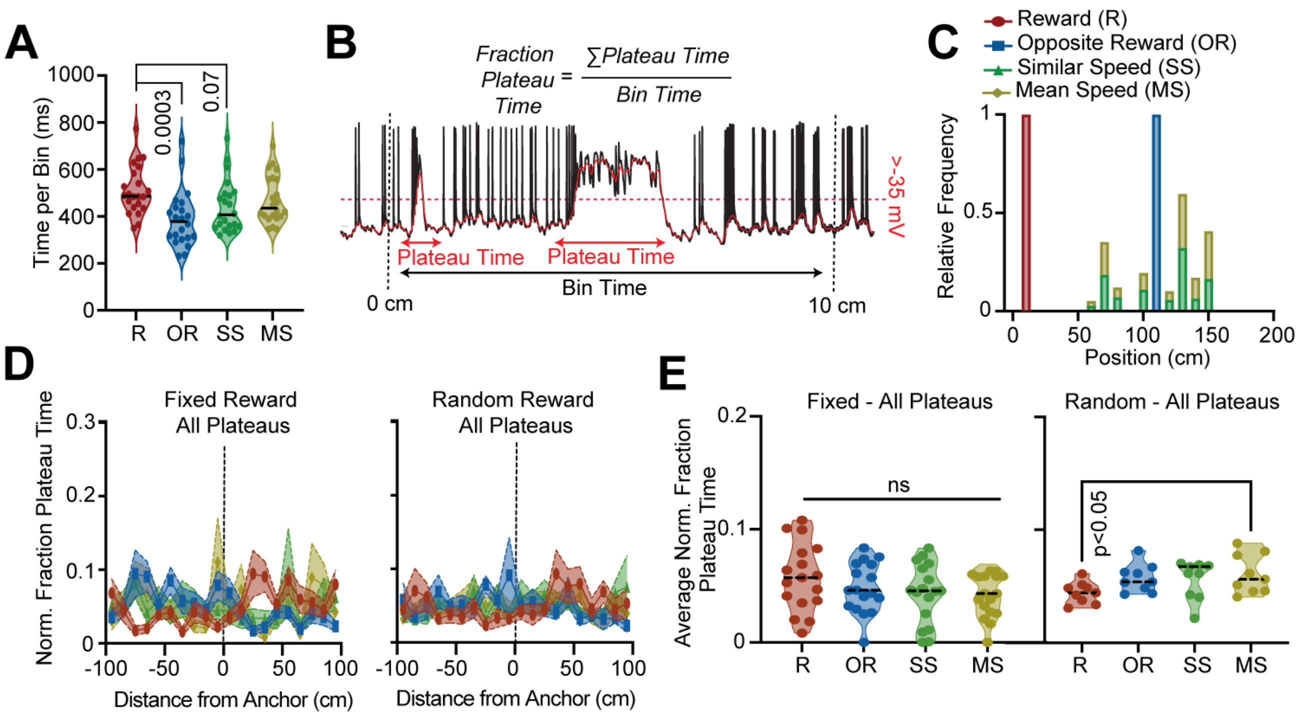


**Supplemental Figure 3: Subiculum plateaus are associated with the reward location.** (A) Average time spent ±40 cm around each anchor (R reward, OR opposite reward location, SS similar speed, MS mean speed). (B) Illustration of fraction plateau time calculation: plateau time per spatial bin divided by dwell time in that bin. (C) Histogram showing anchor locations. (D) Fraction plateau time for all anchors (see C for legend) for all plateaus in fixed (left) and random reward conditions (right). (E) Quantification of the data shown in (D). All line plots are plotted as mean±SEM, unless noted otherwise. Detailed statistical information is listed in Supplementary Table 1.

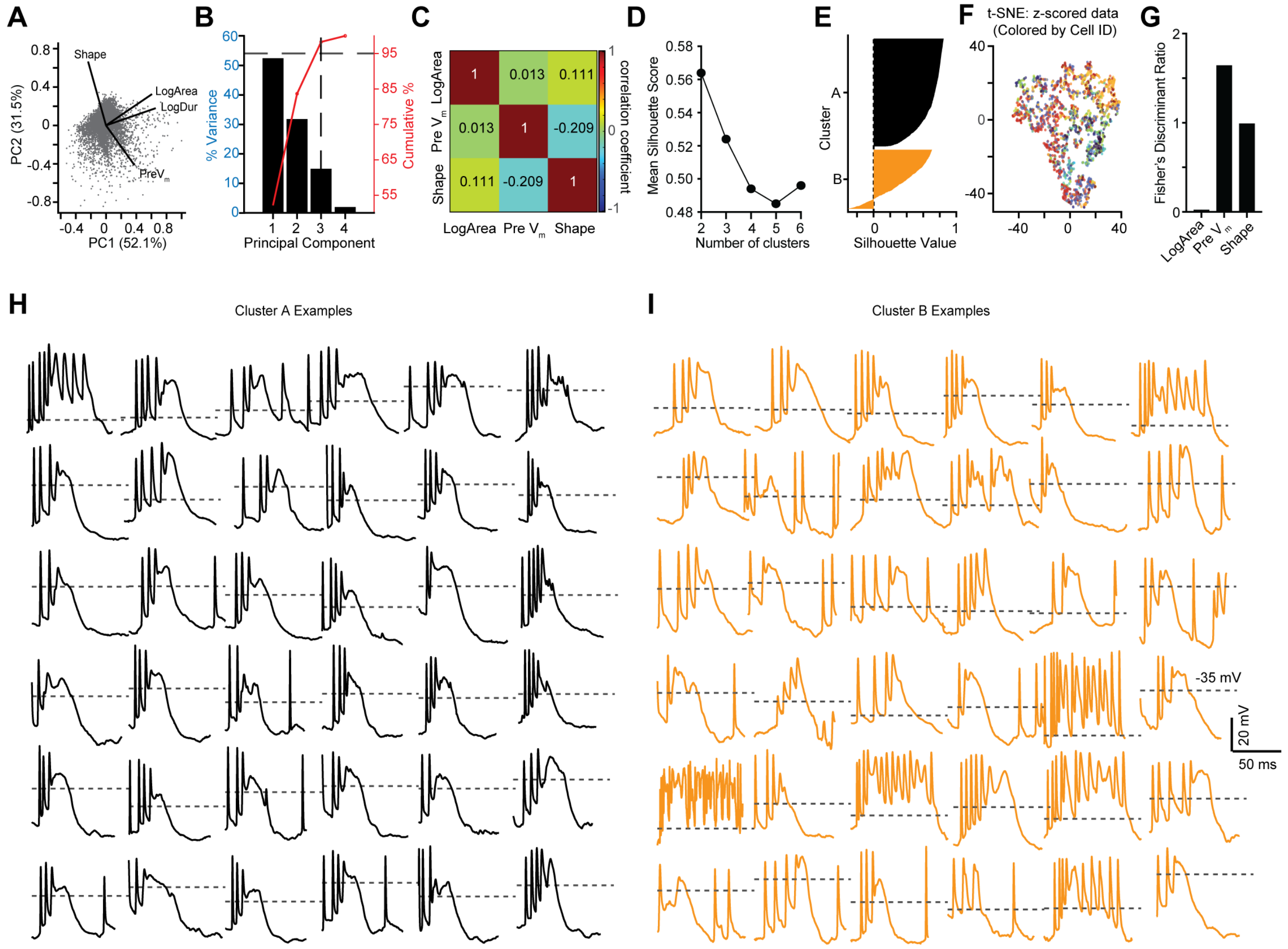


**Supplemental Figure 4: Validation and low-dimensional structure of unsupervised clustering of plateaus.** (A) PCA biplot displaying data points projected onto the first two principal components (PC1 and PC2), which account for 52.1% and 31.5% of the variance, respectively. Vectors represent the loadings for the features Shape, LogArea, LogDuration, and Pre $V_m$. (B) Percentage of variance explained by each principal component (black bars, left axis) and the cumulative percentage of variance (red line, right axis). The dashed lines indicate that the first three components capture ~95% of the total variance. (C) Correlation matrix showing pairwise correlations between the three parameters used for k-means clustering (LogArea, Pre $V_m$, shape). (D) Mean Silhouette Score analysis for cluster counts ranging from 2 to 6. (E) Silhouette plot illustrating the consistency of the data within the two identified clusters. The width of the silhouette indicates the degree of separation and cohesion for each point within its cluster. (F) t-SNE visualization of plateau events following k-means clustering, colored by cell identity. (G) Fisher's Discriminant Ratio for the three parameters used in the k-means clustering. (H) Raw $V_m$ examples of cluster A plateaus. Dotted lines indicate -35 mV. (I) same as (H) but for cluster B plateaus.

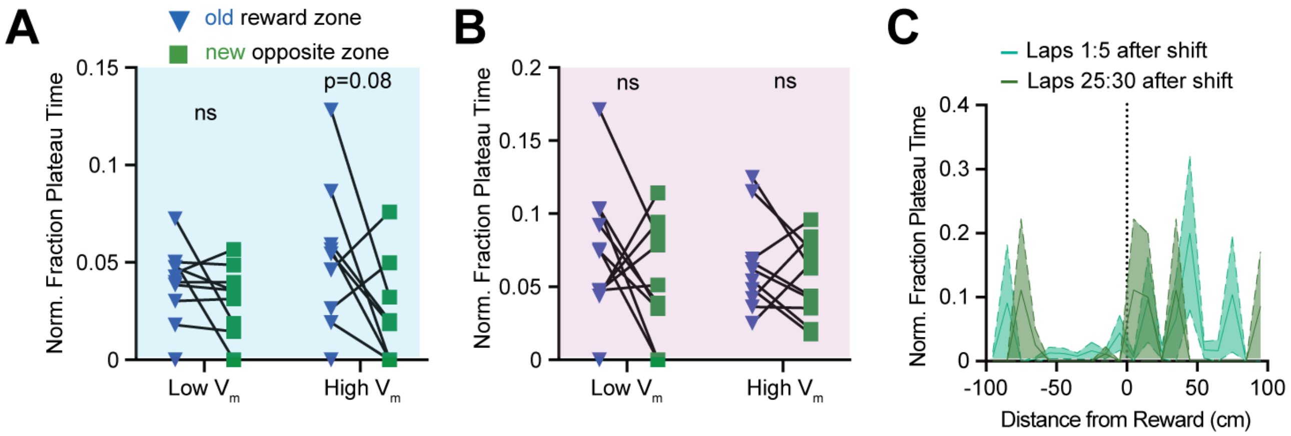


**Supplemental Figure 5: Quantification of plateaus at the original reward location** (A) Comparison between anticipatory plateaus in the original reward zone (learning control) and the new opposite zone (memory zone). (B) same as (A) but for the consumption region. (C) The change in the fraction plateau time of the high $V_m$ plateaus in the first 5 laps after reward shift (cyan) versus laps 25-30 after reward shift (green). Detailed statistical information is listed in Supplementary Table 1.

Supplemental Table 1. Statistical Comparisons

| Figure / Panel | Comparison / Measure | Statistical Test | Test Statistic | p-value |
|---|---|---|---|---|
| **Fig.4E** | Low vs High $V_m$ plateau width | Mann–Whitney U | Median Low $V_m$ Plateaus = 25.15 ms; High $V_m$ Plateaus = 30.55 ms | <0.0001 |
| **Fig.4F** | Spike rate before Low vs High $V_m$ plateaus | Mann–Whitney U | Median Low $V_m$ Plateaus = 15 Hz; High $V_m$ Plateaus = 25 Hz | <0.0001 |
| **Fig.4K (Low $V_m$ Plateaus)** | Clustering around Anchors (Fixed Reward) | Kruskal–Wallis | H(4)=17.19 | 0.0018 |
| **Post-hoc tests: Reward vs. Opposite Reward** | | Dunn's test | Mean Rank difference: 28.65 | 0.0005 |
| **Reward vs. Similar Speed** | | | 22.42 | 0.0075 |
| **Reward vs. Exact Speed** | | | 20.53 | 0.0129 |
| **Reward vs. Stop>Run** | | | 30.61 | 0.0003 |
| **Fig.4K (High $V_m$ Plateaus)** | Clustering around Anchors (Fixed Reward) | Kruskal–Wallis | H(4)=3.04 | 0.551 |
| **Fig.4N (Low $V_m$ Plateaus)** | Clustering around Anchors (Random Reward) | Kruskal–Wallis | H(4)=1.963 | 0.7427 |
| **Fig.4N (High $V_m$ Plateaus)** | Clustering around Anchors (Random Reward) | Kruskal–Wallis | H(4)=11.31 | 0.0233 |
| **Post-hoc tests: Reward vs. Opposite Reward** | | Dunn's test | Mean Rank difference: 16.67 | 0.0071 |

| | | | | |
|---|---|---|---|---|
| **Reward vs. Similar Speed**<br><br>**Reward vs. Exact Speed**<br><br>**Reward vs. Stop>Run** | | | 12.44<br><br>5.722<br><br>0.4444 | 0.0444<br><br>0.3552<br><br>0.9428 |
| **Fig.4Q** | Subthreshold $V_m$ across anchors (Fixed Reward) | Kruskal–Wallis | H(3)=0.063 | 0.99 |
| **Fig.4R** | Normalized firing rate across anchors (Fixed Reward) | Kruskal–Wallis | H(3)=2.1 | 0.553 |
| **Fig. 5D** | Cluster A vs B pre $V_m$ | Mann–Whitney U | Median A=-49.14 mV; B=-44.35 mV | <0.0001 |
| **Fig. 5E** | Cluster A vs B Area | Mann–Whitney U | Median A=329.4 ms*mV; B=281.6 ms*mV | <0.0001 |
| **Fig. 5F** | Cluster A vs B Duration | Mann–Whitney U | Median A=26.25 ms; B=26.7 ms | 0.06 |
| **Fig. 5G** | Cluster A vs B Pearson correlation coefficient | Mann–Whitney U | Median A=0.71; B=0.52 | <0.0001 |
| **Fig. 5J (Cluster A plateaus)**<br><br>**Post-hoc tests: Reward vs. Opposite Reward**<br><br>**Reward vs. Similar Speed**<br><br>**Reward vs. Exact Speed**<br><br>**Reward vs. Stop>Run** | Clustering around Anchors (Fixed Reward) | Kruskal–Wallis<br><br>Dunn's test | H(4)=30.76<br><br>Mean Rank difference: 54.14<br><br>39.86<br><br>41.75<br><br>47.46 | <0.0001<br><br><0.0001<br><br>0.0009<br><br>0.0005<br><br><0.0001 |

| | | | | |
|---|---|---|---|---|
| **Fig. 5K (Cluster B plateaus)** | Clustering around Anchors (Fixed Reward) | Kruskal–Wallis | H(4)=3.396 | 0.4938 |
| **Fig.6G (Learning)** | Control vs Learning (consumption) | Wilcoxon signed-rank | Low $V_m$ Plateaus: W=19; High $V_m$ Plateaus: W=29 | 0.375; 0.160 |
| **Fig.6G (Memory)** | Control vs Memory (consumption) | Wilcoxon signed-rank | Low $V_m$ Plateaus: W=40; High $V_m$ Plateaus: W=12 | 0.083; 0.64 |
| **Fig.6J (Learning)** | Control vs Learning (anticipatory) | Wilcoxon signed-rank | Low $V_m$ Plateaus: W=55; High $V_m$ Plateaus: W=7 | 0.002; 0.769 |
| **Fig.6J (Memory)** | Control vs Memory (anticipatory) | Wilcoxon signed-rank | Low $V_m$ Plateaus: W=29; High $V_m$ Plateaus: W=48 | 0.16; 0.032 |
| **Supp. Fig. 3A** | Time spent across anchors | Kruskal–Wallis | H(3)=16.03 | 0.001 |
| **Supp. Fig. 3E (Fixed Reward)** | Fraction plateau time | Kruskal–Wallis | H(3)=2.74 | 0.434 |
| **Supp. Fig. 3E (Random Reward)** | Fraction plateau time | Kruskal–Wallis | H(3)=4.73 | 0.19 |
| **Supp. Fig. 5A** | Anticipatory learning vs memory zones | Wilcoxon signed-rank | Low $V_m$ Plateaus: W=41; High $V_m$ Plateaus: W=26 | 0.129; 0.078 |
| **Supp. Fig. 5B** | Consumption learning vs memory zones | Wilcoxon signed-rank | Low $V_m$ Plateaus: W=27; High $V_m$ Plateaus: W=13 | 0.19; 0.56 |
